# Engineering oleaginous yeast *Yarrowia lipolytica* for violacein production: extraction, quantitative measurement and culture optimization

**DOI:** 10.1101/687012

**Authors:** Yingjia Tong, Jingwen Zhou, Liang Zhang, Peng Xu

**Author notes:** Corresponding author. E-mail address (PX) and (LZ).

## Abstract

Violacein is a naturally occurring anticancer therapeutic with deep purple color. Yeast fermentation represents an alternative approach to efficiently manufacturing violacein from inexpensive feedstocks. In this work, we optimized the extraction protocol to improve violacein recovery ratio and purity from yeast culture, including the variations of organic solvents, the choice of mechanical shear stress, incubation time and the use of cell wall-degrading enzymes. We also established the quantitative correlation between HPLC and microplate reader method. We demonstrated that both HPLC and microplate reader are technically equivalent to measure violacein from yeast culture. Furthermore, we optimized the yeast cultivation conditions, including carbon/nitrogen ratio and pH conditions. Our results indicated that ethyl acetate is the best extraction solvent with glass beads grinding the cell pellets, the maximum violacein and deoxyviolacein production was 70.04 mg/L and 5.28 mg/L in shake flasks, respectively. Violacein purity reaches 86.92% at C/N ratio of 60, with addition of 10 g/L CaCO_3_ to control the media pH. Taken together, the development of efficient extraction protocol, quantitative correlation between HPLC and microplate reader, and the optimization of culture conditions set a new stage for engineering violacein production in *Y. lipolytica*. This information should be valuable for us to build a renewable and scalable violacein production platform from the novel host oleaginous yeast species.

## Introduction

Violacein and deoxyviolacein belong to bisindol pigments with deep purple color, which are derived from tryptophan biosynthetic pathway and naturally produced by a number of marine bacteria such as *Janthinobacterium lividum* [1, 2], *Chromobacterium violaceum* [3–5], *Pseudoalteromonas luteoviolacea* [6], *et al.* Clinical trials and biomedical studies indicate both compounds possess strong antibacterial [7, 8], anticancer [9], antiviral [10], trypanocidal [11] and antiprotozoal [12] properties. These characteristics make violacein a superior chemical scaffold and drug candidate for the development of clinically active agents.

Violacein biosynthetic pathway was first discovered by Pemberton et al. [13] in 1991 and was fully characterized by Balibar et al. [14] and Sanchez et al. [15] in 2006. Branched from the L-tryptophan pathway, the violacein biosynthetic pathway involves five steps, encoded by *vio*A, *vio*B, *vio*C, *vio*D and *vio*E [16], which were organized in an operon form containing all the five genes. Two molecules of L-tryptophan are oxidatively condensed by vioA and vioB to form indole-3-pyruvic acid (IPA). Then, IPA is decarboxylated to form protodeoxyviolaceinic acid via vioE. Subsequently, protodeoxyviolaceinate is reduced to violacein by vioD and VioC. Without involving the first reduction step of vioD, deoxyviolacein is formed as the major byproduct [16–18].

At present, most of the reported violacein-producing host are gram-negative bacteria with human pathogenicity. For example, both *C. violaceum* [3–5], and *J. lividum* [1, 2] have been related with serious skin infection in immune-compromised people, and both strain have been classified as biosafety level II bacteria. Although the native host represents some advantage to produce violacein, the pathogenicity significantly limited their industrial application. *Yarrowia lipolytica* has been considered to be non-pathogenic and has been classified as ‘generally regarded as safe’ (GRAS) by the US Food and Drug Administration (FDA). For example, *Y. lipolytica* has been widely adopted as host for production of citric acid [19, 20], β-carotenoids[21–23] and β-ionone [24] in food industry. Both *Y. lipolytica* and *C. violacein* were isolated from marine environment with high GC content (up to 65% in the coding sequence), we argue that *Y. lipolytica* might be engineered as a novel production platform for violacein production due to its GRAS status. Different from Baker’s yeast, *Y. lipolytica* is abundant in acetyl-CoA due to its cytosolic ATP citrate lyase (ACL) [25]. *Y. lipolytica* has been reported to grow on a wide range of inexpensive raw materials[26], including glucose, glycerol, xylose[27, 28], volatile fatty acids[29, 30], alcohols and wax alkanes *et al.*

In our previous report[31], the five-gene pathway of violacein (without codon optimization) was assembled and functionally expressed in *Y. lipolytica* using the self-replicating YaliBrick vectors. In this work, we optimized the extraction protocol to isolate violacein from yeast culture, and developed the quantitative correlation between HPLC and microplate reader method. We demonstrated that both HPLC and microplate reader are technically equivalent to measure violacein from yeast culture, especially, the microplate reader method has the advantage for high throughput screening. Furthermore, we optimized the yeast cultivation conditions, including carbon/nitrogen ratio and pH conditions. Our results indicated that ethyl acetate is the best extraction solvent, the maximum production of violacein and deoxyviolacein was 70.04 mg/L and 5.28 mg/L in shake flasks, respectively. Violacein purity reaches 86.92% at C/N ratio of 60, with addition of 10 g/L CaCO_3_ to control the media pH. Taken together, the development of extraction protocol, quantitative correlation between HPLC and microplate reader, and the optimization of culture conditions set a new stage for engineering violacein production in *Y. lipolytica*. This information should be valuable for us to build a renewable and scalable violacein production platform from the novel host oleaginous yeast species.

## 2. Materials and methods

### 2.1. Strains and culture conditions

The auxotrophic *Y. lipolytica* po1g (Leu−) was purchased from Yeastern Biotech Company (Taipei, Taiwan). *Y. lipolytica* XP1 harboring the plasmid pYaliA1-vioDCBAE was stored in our lab [31]. YNB medium (C/N=60, 80, 100, 120) contains 1.7 g/L yeast nitrogen base (without amino acids and ammonium sulfate) (Difco), 1.1 g/L ammonium sulfate (Sigma-Aldrich), 0.69 g/L CSM-Leu (Sunrise Science Products, Inc.), and 30, 40, 50, 60 g/L glucose, respectively. YNB medium (pH 6.0, 6.5, 7.0, 7.5) contains 1.7 g/L yeast nitrogen base (without amino acids and ammonium sulfate) (Difco), 1.1 g/L ammonium sulfate (Sigma-Aldrich), 0.69 g/L CSM-Leu (Sunrise Science Products, Inc.), 30 g/L glucose, and was adjusted to pH 6.0, 6.5, 7.0, 7.5, respectively, through Na_2_HPO_4_ and NaH_2_PO_4_. YNB medium with CaCO_3_ was made with YNB media supplemented with 10 g/L CaCO_3_. Selective YNB plates were made with YNB media supplemented with 20 g/L Bacto agar (Difco).

### 2.2 Seeds and fermentation procedure

For preparing seed inocula, strains were cultivated into 50 mL CSM-Leu medium containing 20 g/L glucose, 5.0 g/L ammonium sulfate (Sigma-Aldrich), 0.69 g/L CSM-Leu (Sunrise Science Products, Inc.) and 1.7 g/L yeast nitrogen base (without amino acids and ammonium sulfate) (Difco) in 250 mL shake flasks for 24 h with agitation 250 rpm at 30 °C. For C/N ratio and pH optimization in shake flask, 5% (v/v) of the seed culture was inoculated to the corresponding medium, with different C/N ratio, pH and CaCO_3_. All the strains were cultured at 250 rpm, 30 °C up to 120 hours for violacein production.

### 2.3 Analytical methods

Violacein standard was purchased from Sigma-Aldrich and was dissolved in methanol. The maximum absorption wavelength of violacein was determined through full wavelength scanning from 230 nm to 750 nm by microplate reader (Synergy H1 Hybrid Multi-Mode Microplate Reader, BioTek, Winooski, VT).

A standard curve for violacein was prepared by diluting the standard (Sigma-Aldrich) with methanol into concentration of 10, 20, 30, 40, 50, 60, 70, 80, 90 and 100 mg/L. The analysis of violacein were performed on high-performance liquid chromatography (HPLC) (Agilent Technologies 1220 Infinity II LC, USA) using Agilent Eclipse Plus C18 column (4.6*100 mm, 3.5 μm, USA.), equipped with UV detector at 570 nm. Mobile phase A was H_2_O with 0.1% acetate acid, and mobile phase B was methanol with 0.1% acetate acid. A stepwise gradient elution at a flow rate of 0.4 mL/min was used, following the gradient profiles as below: ramping from 100% to 20% A in 0~5 min, maintaining 20% A (80% B) in 5~8 min, ramping from 20% to 100% A in 8~12 min, and maintaining100% A in 12~15 min. Deoxyviolacein standard was diluted with methanol into concentrations at 0.6, 1, 1.8, 2.5, 3, 3.5, and 4 mg/L, and was analyzed by HPLC with the same method of violacein. A spectrophotometer-based standard curve was also developed through microplate reader at 570 nm. The violacein standard was diluted with ethanol at the concentration of 2, 4, 6, 8, 10 mg/L.

The biomass was measured as dry cell weight. 500 μL of the yeast cells were centrifuged at 13,000 rpm for 5 minutes. The upper broth was discarded and the cell pellet were dried in an oven at 60 °C until the weight did not change, then calculate the difference in the weight of microcentrifuge tube.

### 2.4 Violacein extraction methods

During optimization of the violacein extraction, 500 μL of the yeast culture was harvested after growth of 12 h in CSM-leu media. Methods used to extract violacein are listed in Table 1.

**Table 1.** Methods used for violacein extraction in this work.

| Nu<br>mber | Methods |
| --- | --- |
| 1 | Add 500 μL ethyl acetate, incubate at 30 °C for 24 h |
| 2 | Add 500 μL ethyl acetate, vortex with glass beads for 1 min, and incubate at 30 °C for 24 h |
| 3 | Add 500 μL ethyl acetate, vortex with glass beads for 5 min, and incubate at 30 °C for 24 h |
| 4 | Add 500 μL cell lysis buffer, incubate at 25 °C for 2 h, and vortex with glass beads for 2 min |
| 5 | Add 500 μL cell lysis buffer, incubate at 25 °C for 2 h, and vortex with glass beads for 4 min |
| 6 | Add 500 μL methanol, incubate at 30 °C for 24 h |
| 7 | Add 500 μL methanol, vortex with glass beads for 1 min, and incubate at 30 °C for 24 h |
| 8 | Add 500 μL methanol, vortex with glass beads for 5 min, and incubate at 30 °C for 24 h |
| 9 | Add 500 μL methanol, incubate at 75 $^{\circ}$C for 30 min |
| 10 | Add 500 μL methanol, vortex with glass beads for 1 min and incubate at 75 $^{\circ}$C for 30 min |

After extraction, the cell lysate were centrifuged at 13,000 rpm for 5 min. The upper phase containing violacein was collected and transformed into another centrifuge tube and analyzed through HPLC and microplate reader. These extraction procedures were repeated until all the cells became pale.

## 3 Results and discussion

### 3.1 Comparison of violacein standard curve with HPLC and plate reader

To establish a quick method to detect violacein, we decided to compare how microplate reader approach is correlated with HPLC method. With 200 μL of 20, 40, 60, 80, and 100 mg/L violacein standards, we performed full wavelength scanning from 230 nm to 750 nm. As shown in Fig. 1, the maximum absorption wavelength of all the samples were 570 nm, which was consistent with the maximum absorption wavelength previously reported [32]. As the concentration of the violacein standard increases, the absorbance increases, following the Lambert–Beer law. For this reason, 570 nm is used for subsequent HPLC and microplate reader detection.

**Fig. 1.**
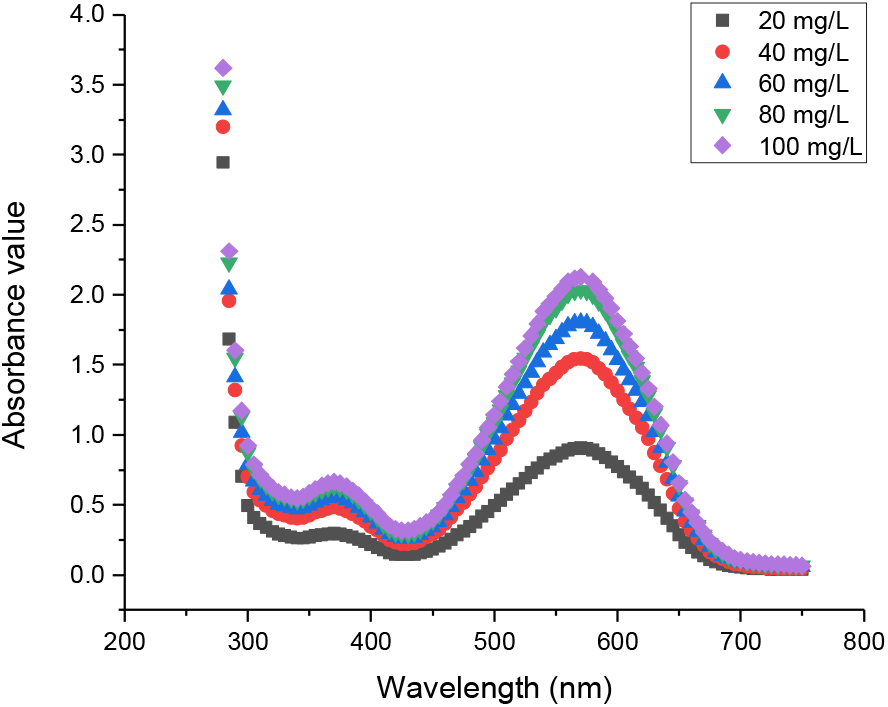
UV-Visible absorption spectrum of violacein standards.

To establish the HPLC standard curve, we prepared violacein standards from 10 mg/L to 100 mg/L, diluted with methanol. As shown in Fig. 2, all the violacein samples have the same retention time (10.6 min), the peak height and area of the violacein standard chromatogram were increased as we increase the violacein concentration. We next constructed the HPLC calibration curve and determined thecorrelation coefficient (R^2^) was 0.9956 (Fig. 3), indicating the reliability of HPLC method to detect violacein from 10 mg/L to 100 mg/L. The correlation between peak area and violacein concentration was found to follow this regression equation: y=0.017x-3.2774.

**Fig. 2.**
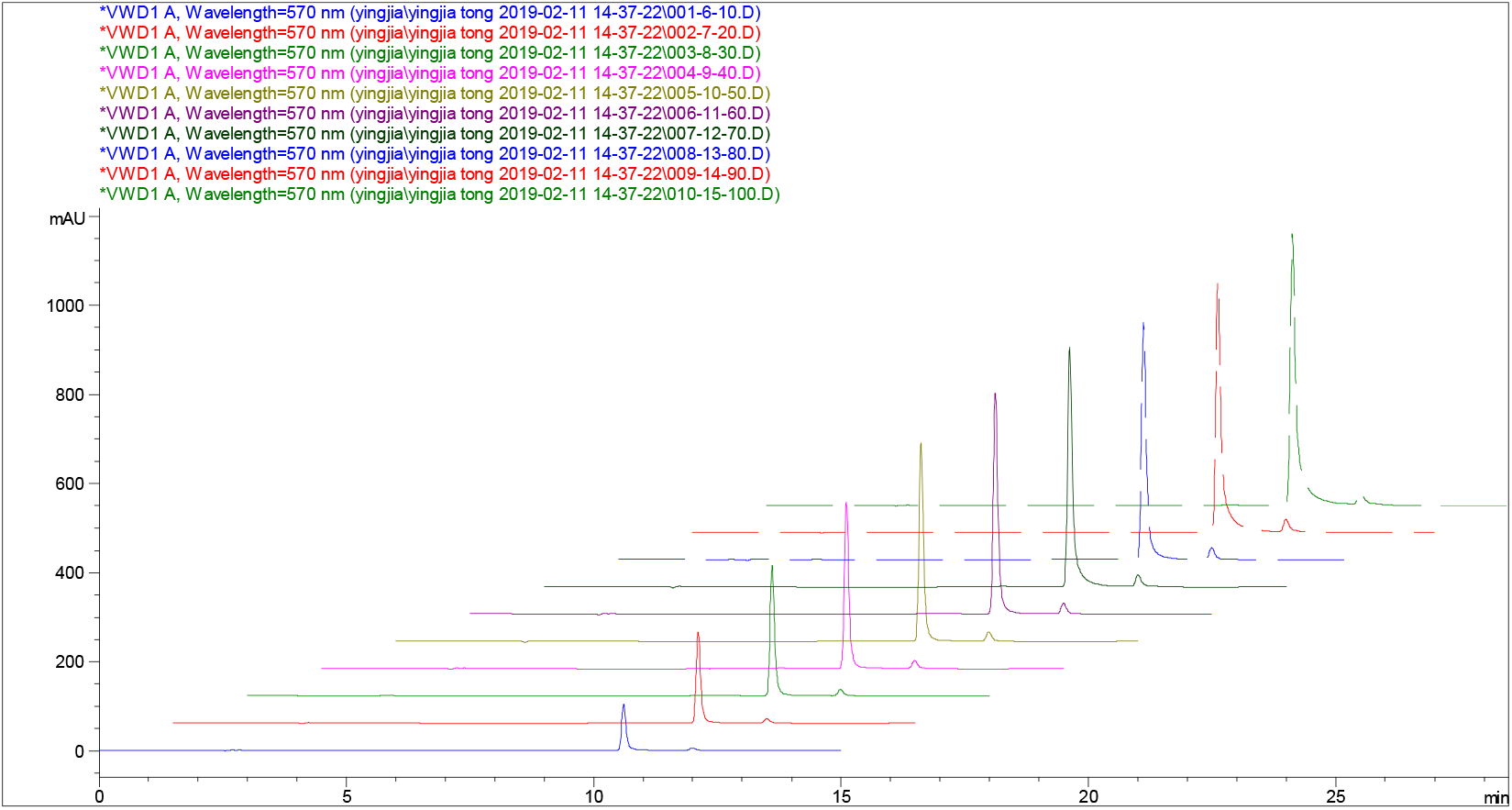
HPLC profile of violacein standards from 10 mg/L to 100 mg/L. HPLC profile were overlayed with the Agilent ChemStation software.

**Fig. 3.**
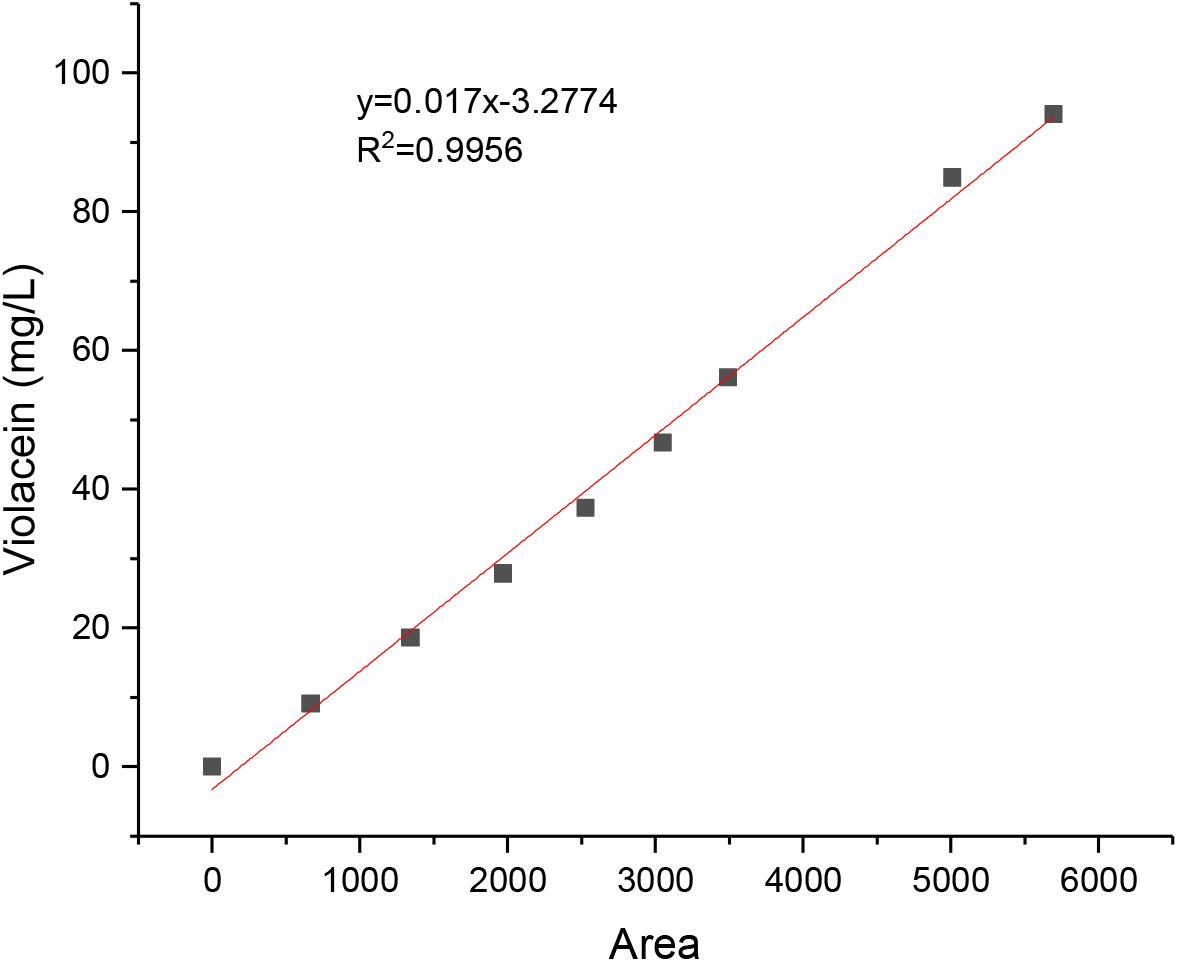
Regression curve between peak area and violacein concentration determined by HPLC at 570 nm.

We next built the violacein calibration curve with microplate reader. Absorbance at 570 nm was recorded with sequentially diluted violacein standards. We found the correlation coefficients (R^2^) of plate reader method is about 0.9973 (Fig. 4), which is equivalent to HPLC method (Fig. 3) in terms of accuracy and reliability. The correlation between absorbance and violacein concentration was found to follow this regression equation: y=15.699x-0.1938.

**Fig. 4.**
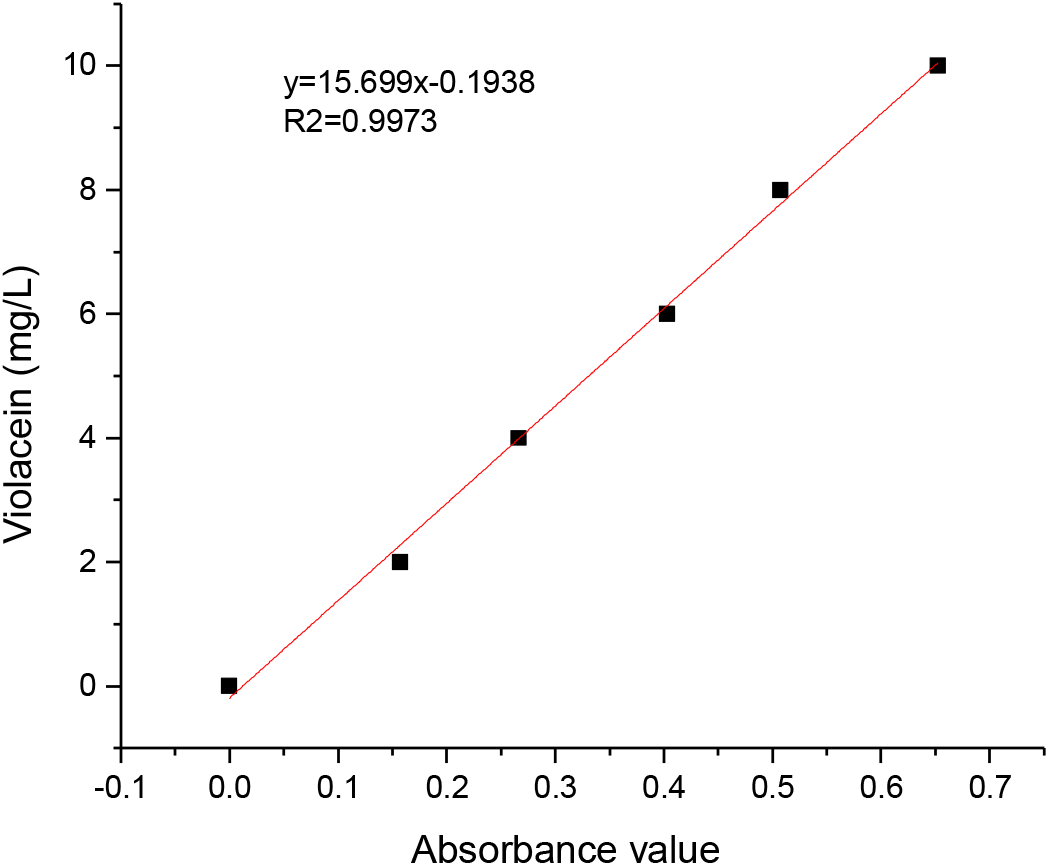
Regression curve between absorbance and violacein concentration determined by microplate reader at 570 nm.

The deoxyviolacein calibration curve was also determined by HPLC chromatogram with the correlation coefficients (R^2^) about 0.9962 (Fig. 5). The correlation between peak area and deoxyviolacein concentration was found to follow this regression equation: y=0.0157x-0.0821.

**Fig. 5.**
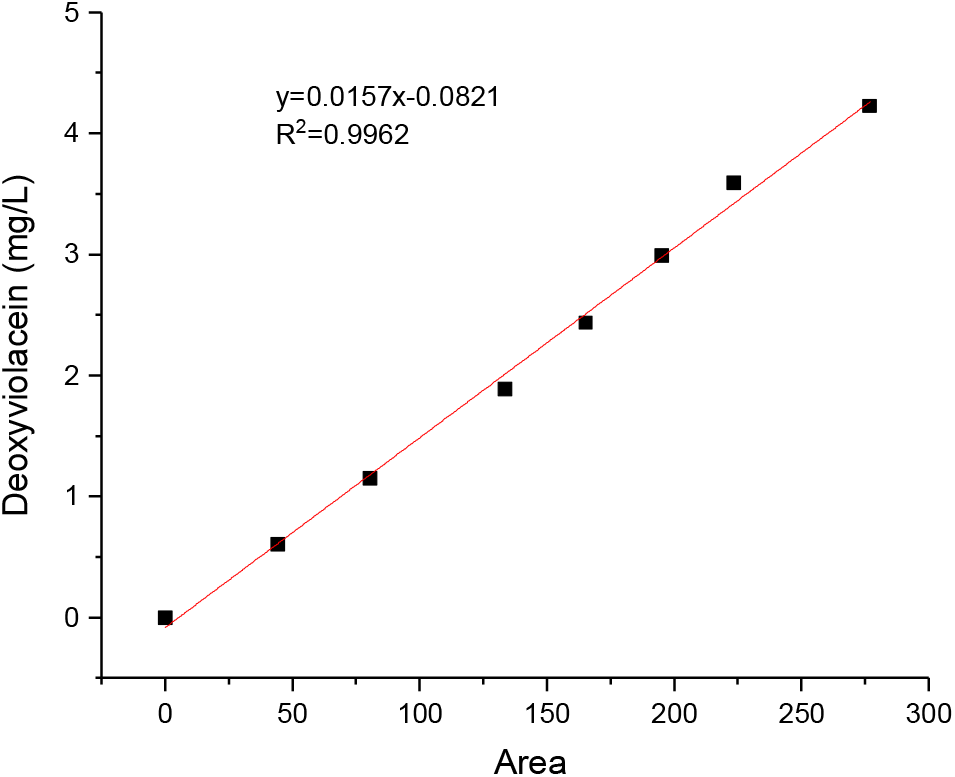
Regression curve between peak area and deoxyviolacein concentration determined by HPLC at 570 nm.

It should be emphasized that all the regression curves in Fig. 3, Fig. 4 and Fig. 5 demonstrated great linear relationship between peak area/absorbance value and concentration, indicating the accuracy and reliability of the reported method. In contrast to the tedious HPLC method, the plate reader methods potentially could be used as a standard protocol for high/medium throughput screening of large library of strains. Both the HPLC and plate reader approach should be considered technically equivalent, if strains or colonies already demonstrate qualitatively purple color.

### 3.2 Violacein extraction method optimization

Different from *E. coli*, *Y. lipolytica* is a dimorphic yeast surrounded by a rigid cell wall with chitin-like polymer consisting of mannose and galactose [33]. This cell wall protects the cell from osmotic or mechanic pressure and maintains cell shape. We observed that most of the violacein were trapped intracellularly. Most of the previously reported violacein extraction protocol is based on *E. coli* culture [18, 34]. It is important to develop efficient extraction protocol, so that we can release intracellular violacein from *Y. lipolytica*. We investigated a number of cell disruption and extraction method to maximize violacein recovery ratio and purity. Based on previous report [18, 34], we have considered solvent type (ethyl acetate or methanol), mechanical shear stress (glass beads and vortex) and enzyme digestion in our tested methods (Table 2), which are commonly accessible to biological labs. We specifically evaluated 10 different methods to extract violacein from 500 μL yeast culture. HPLC grade ethyl acetate and methanol with different shear stress settings were used to extract violacein (Table 2). To break up cell wall, cell lysis enzyme Zymolyase 20-T were also tested (Table 2).

**Table 2.** Methods used for violacein extraction and the resulting violacein yield and purity. To save time, all sample were taken at 12 hour of cultivation.

| Number | Methods | Yield (mg/L) | Violacein Purity |
| --- | --- | --- | --- |
| 1 | Add 500 μL ethyl acetate, incubate at 30 $^{\circ}$C for 24 h | 14.99 | 88.41% |
| 2 | Add 500 μL ethyl acetate, vortex with glass beads for 1 min, and incubate at 30 $^{\circ}$C for 24 h | 14.77 | 86.28% |
| 3 | Add 500 μL ethyl acetate, vortex with glass beads for 5 min, and incubate at 30 $^{\circ}$C for 24 h | 15.76 | 86.97% |
| 4 | Add 500 μL Zymolyase 20-T, incubate at 25 $^{\circ}$C for 2 h, and vortex with glass beads for 2 min | 8.12 | 69.28% |
| 5 | Add 500 μL Zymolyase 20-T, incubate at 25 $^{\circ}$C for 2 h, and vortex with glass beads for 4 min | 8.48 | 68.25% |
| 6 | Add 500 μL methanol, incubate at 30 $^{\circ}$C for 24 h | 8.99 | 80.45% |
| 7 | Add 500 μL methanol, vortex with glass beads for 1 min, and incubate at 30 $^{\circ}$C for 24 h | 11.43 | 79.44% |
| 8 | Add 500 μL methanol, vortex with glass beads for 5 min, and incubate at 30 $^{\circ}$C for 24 h | 10.47 | 79.01% |
| 9 | Add 500 μL methanol, incubate at 75 $^{\circ}$C for 30 min | 7.87 | 78.63% |
| 10 | Add 500 μL methanol, vortex with glass beads for 1 min and incubate at 75 $^{\circ}$C for 30 min | 11.26 | 77.07% |

All the cultivated colonies demonstrated cyanish color and the extractions showed a deep purple color (Fig. 6). After HPLC determination, the retention times (10.6 min) of all the extractions were consistent with the violacein standard (Fig. 7). As shown in Table 2, violacein yield and purity varied significantly with different extraction methods. Ethyl acetate with glass bead shear stress led to the best extraction efficiency and purity. After incubated at 30 °C for 24 h with 500 μL ethyl acetate, the yield and purity of violacein reaches 14.99 mg/L and 88.24% from 500 μL yeast culture (12 hour of fermentation sample), respectively. Subsequently, despite the slight decline in purity, vortexing with glass beads for 5 min further increased the extraction efficiency by 5.14% to 15.76 mg/L compared to the samples without glass beads. Incubating cells with methanol is also a widely used extraction method [35]. However, we found that the extraction efficiency with methanol was not high. By varying the shear stress and incubation time, the most efficient methanol extraction only yielded 11.43 mg/L violacein, with a purity of 79.44% (Table 2). The different extraction efficiency could be linked with the relatively highly lipophilic nature of violacein. As a result, ethyl acetate demonstrated better extraction efficiency than methanol. At the same time, we also attempted to extract the intracellular violacein by enzymatically digesting the cell wall and breaking cell wall with glass beads. Surprisingly, enzyme digestion led to the lowest extraction efficiency and purity, possibly due to the sequestration of violacein by the lipid bodies in *Y. lipolytica*. Since the yield and purity with zymolase digestion is low, and the enzymatic method is expensive, it is not considered in the subsequent experiments.

**Fig. 6.**
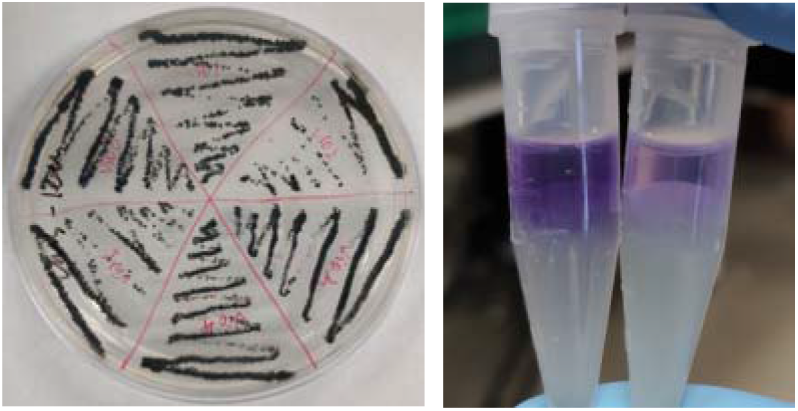
Violacein-producing yeast colony grown on CSM-Leu solid media demonstrated cyanish color (left); and violacein extracted with ethyl acetate from CSM-Leu yeast culture exhibited deep purple color (right).

**Fig. 7.**
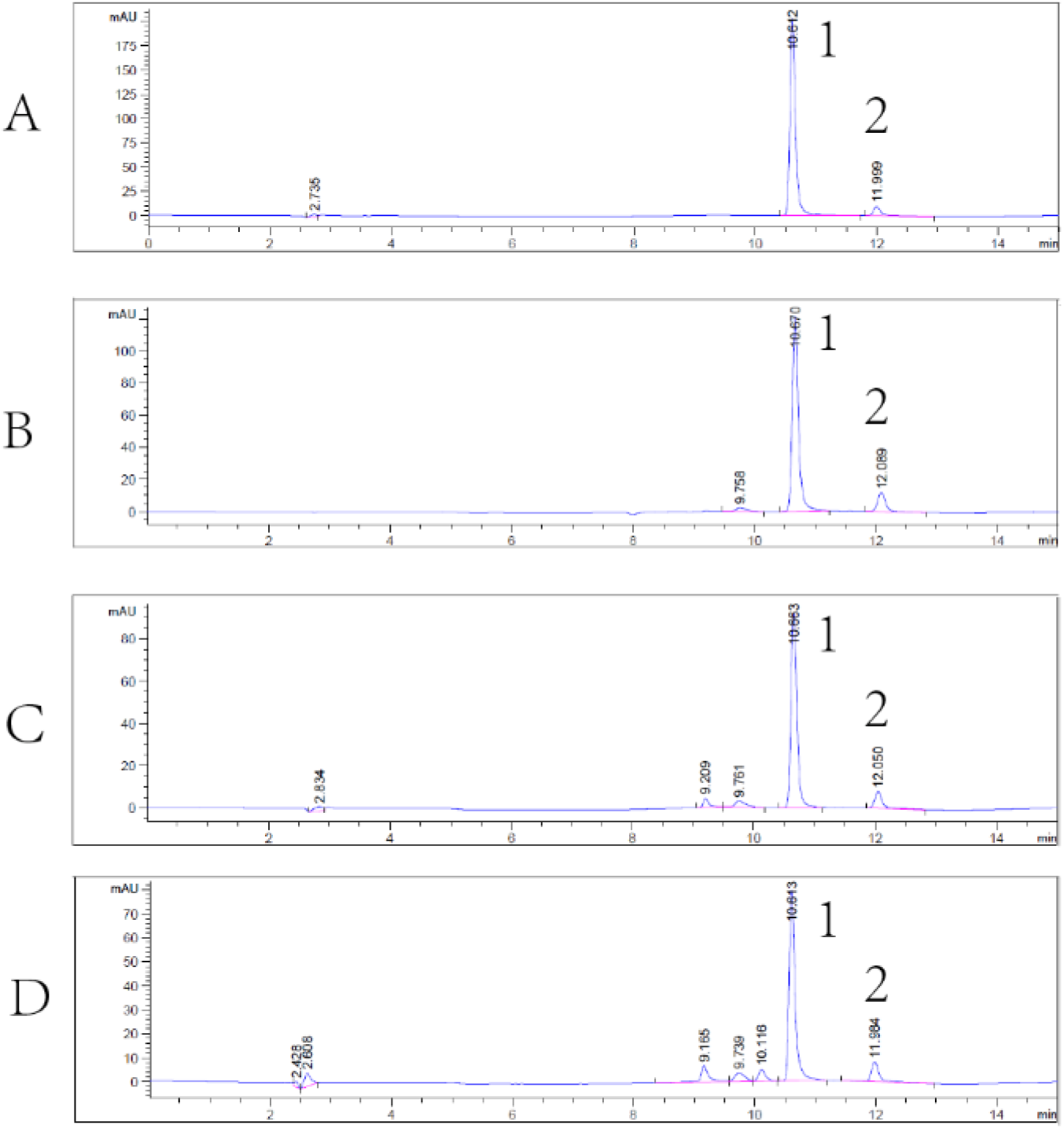
The HPLC profiles of violacein extraction. (A) violacein standard, (B) violacein extracted by ethyl acetate, (C) violacein extracted by methanol, (D) violacein extracted by Zymolyase 20-T. Peaks represented: 1, violacein; 2, deoxyviolacein.

### 3.3 Selection of high violacein-producing colony from shake flask culture

In order to select the yeast colony with highest yield, the laboratory-preserved *Y. lipolytica* XP1 was streaked in CSM-Leu plate at 30 °C. Then, four dark single colonies were tested for shake flask fermentation in CSM-Leu medium with C/N ratio of 80, and incubated at 30 °C for 144 h. As shown in Fig. 7, although all the four biologically replicative colonies were picked from the same plate, HPLC quantification of violacein production varied significantly among these biologically replicative colonies. Colony No. 2 showed the highest yield, reaching 32.75 mg/L of violacein, which was 60% higher than that the colony No. 3 (Fig. 8). This production is consistent with the previously reported result (31.5 mg/L) quantified by microplate reader [31]. The colony-to-colony variations of violacein production may be due to a number of factors, including (1) plasmid copy number variations, (2) genetic integration site variations, or (3) the non-homologous end-joining mechanism of *Y. lipolytica* [36, 37]. Therefore, the colony No. 2 was preserved and selected for subsequent optimization.

**Fig. 8.**
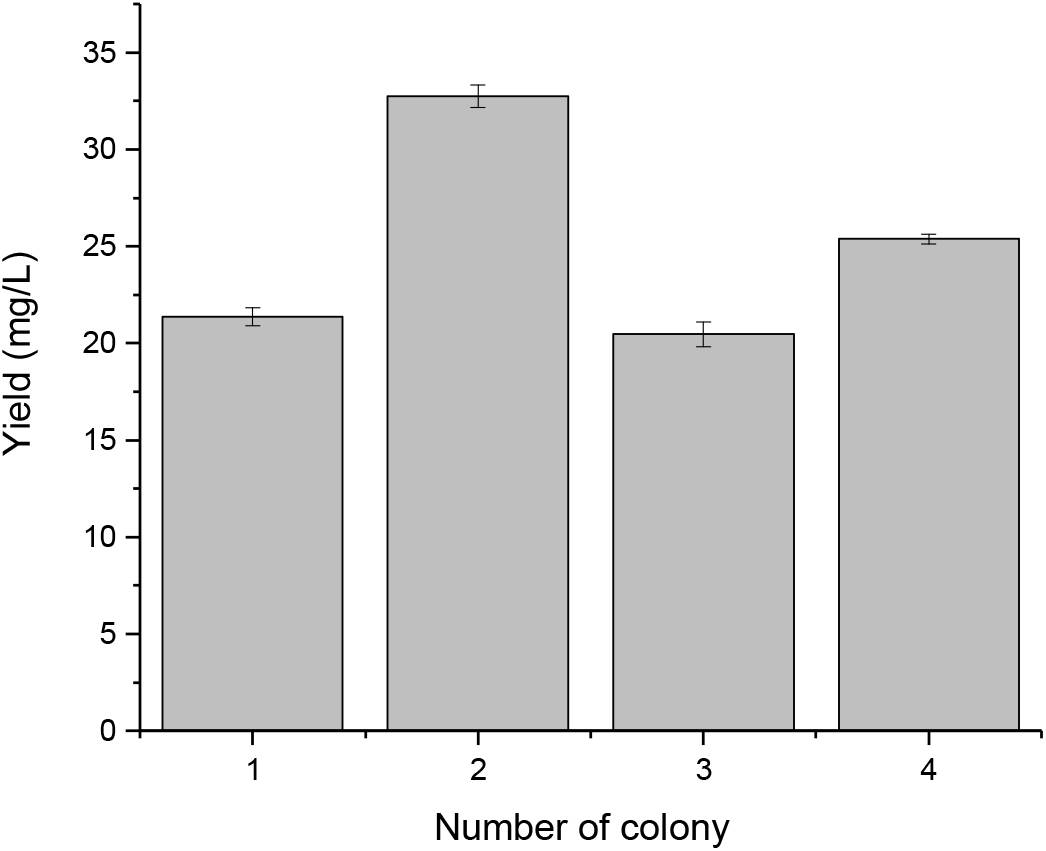
Violacein yield of four biologically replicative colonies cultivated in CSM-Leu media (C/N 80) at 30 °C for 144 hours, yeast culture was extracted with ethyl acetate and violacein were quantified with HPLC.

### 3.4 C/N optimization in shake flask fermentation

Carbon and nitrogen ratio plays a major role in regulating the metabolic activity of lipogenic pathways in *Y. lipolytica* [38]. To understand whether C/N ratio affects vaiolacein production, we tested the violacein production of the recombinant strain *Y. lipolytica* XP1 cultivated in CSM-Leu with C/N ratio fixed at 60, 80,100, and 120, respectively. Since the maximum absorption wavelength of violacein (570 nm) is close to the detection wavelength of the OD_600_, to avoid optical interference, dry cell weight, instead of OD_600_, was used to quantify cell growth. As shown in Fig. 9, the dry cell weight increased as we increased the C/N ratio, possibly due to the increased lipid accumulation. However, violacein and deoxyviolacein production remained unchanged from 38 mg/L to 41 mg/L, regardless of the C/N variations (Fig. 10). Since the yield is slightly increased under the condition of C/N ratio of 60, the subsequent experiment will be carried out under the condition of C/N ratio of 60.

**Fig. 9.**
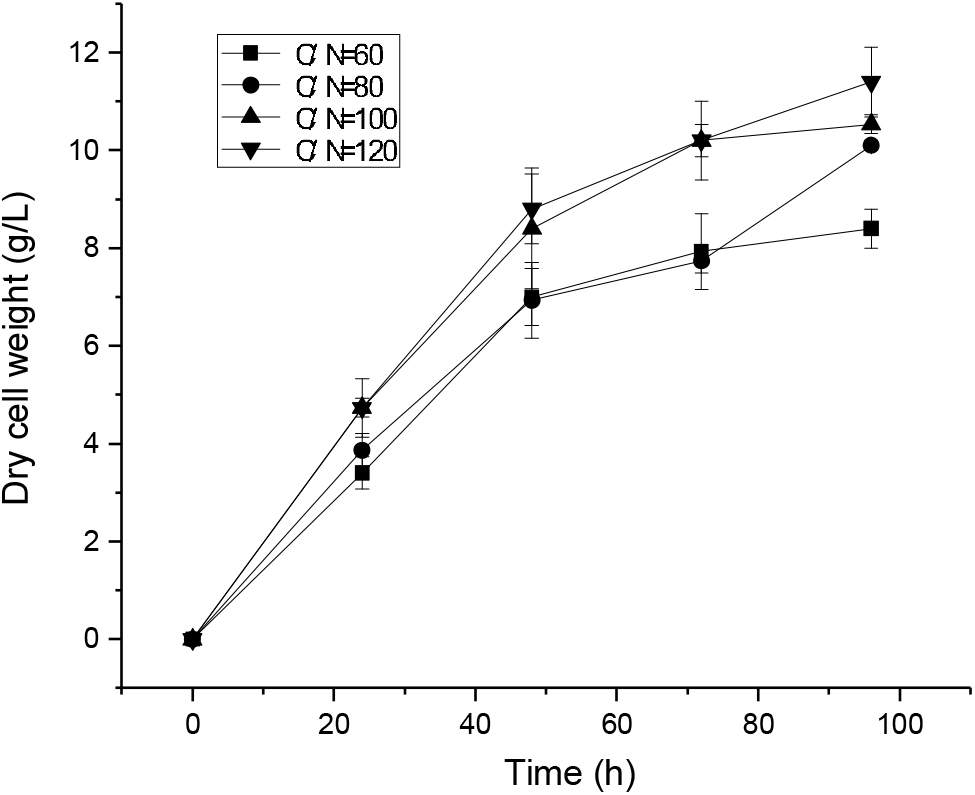
Growth curves of *Y. lipolytica* XP1 in shake flask fermentation with different C/N ratio, cultivated in CSM-leu media at 30 °C.

**Fig. 10.**
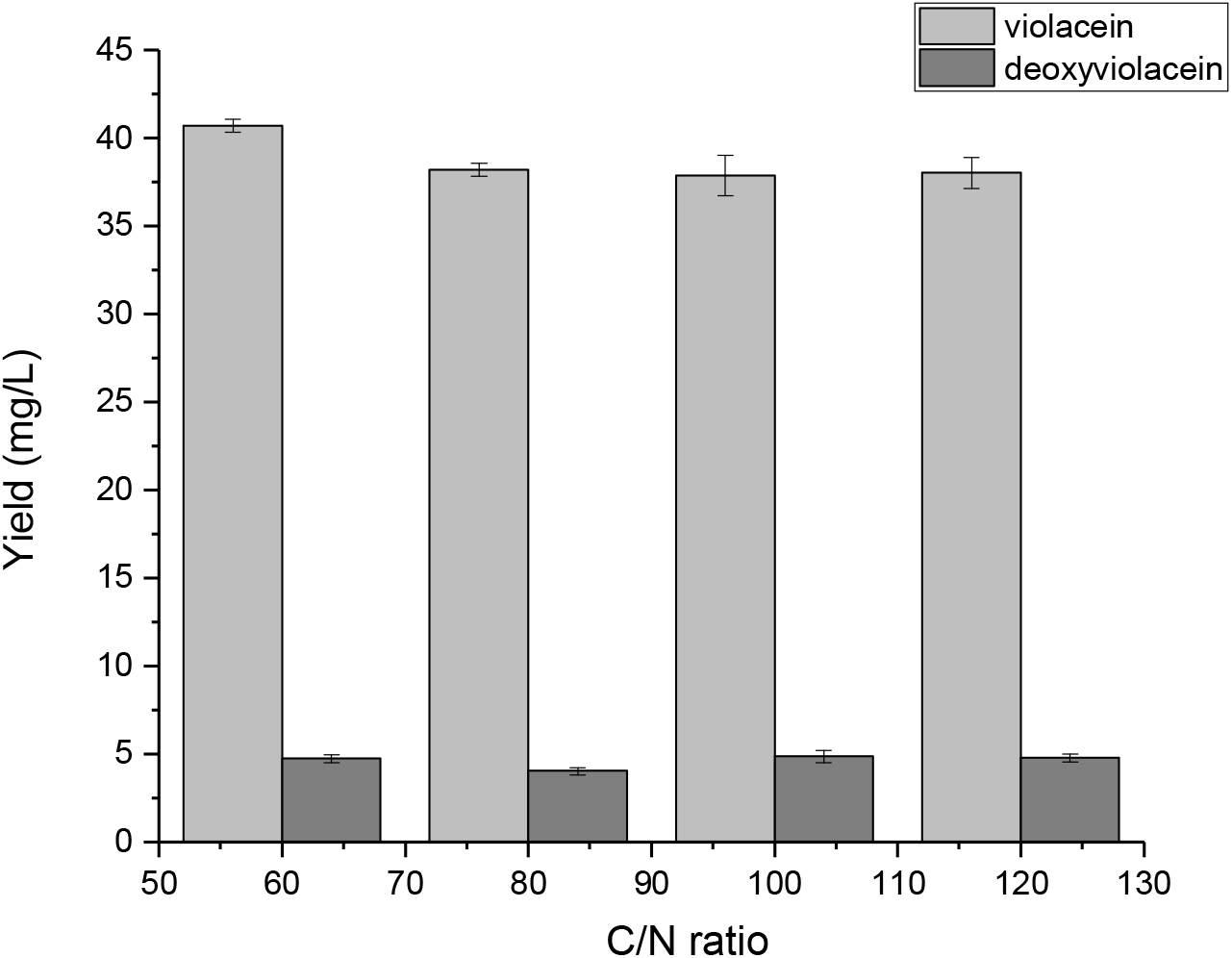
Violacein and deoxyviolacein yield in shake flack fermentation by strain *Y. lipolytica* XP1 with different C/N ratios, cultivated in CSM-leu media at 30 °C. Violacein and deoxyviolacein were quantified by HPLC.

### 3.5 pH optimization in shake flask fermentation

pH has played a major role in regulating cell physiology and metabolic activity. Low pH has been routinely observed in *Y. lipolytica* culture due to the accumulation of organic acids[39]. To evaluate the effects of pH on cell growth and violacein production, the cultivation pH was adjusted by adding 10 g/L CaCO_3_, or by buffering with NaH_2_PO_4_ and Na_2_HPO_4_ to pH 6.0, 6.5, 7.0, and 7.5, with initial unbuffered media as control. As shown in Fig. 11, media pH strongly influences cell growth (data of samples adding 10 g/L CaCO_3_ was not shown, due to insoluble CaCO_3_ crystals interfere with dry cell weight reading). Surprisingly, the yeast grew better under mild acidic or alkaline conditions. For example, when the initial pH is 7.5 or 4.5, the cell growth rate is the fastest, and the dry cell weight reaches 11.13 g/L and 9.54 g/L at 120 hours, respectively. However, when the media pH was maintained at 6.0, 6.5 and 7.0, the cell growth noticeably slowed down, and the dry cell weight was only 7.73, 4.07 and 4.50 g/L, respectively.

**Fig. 11.**
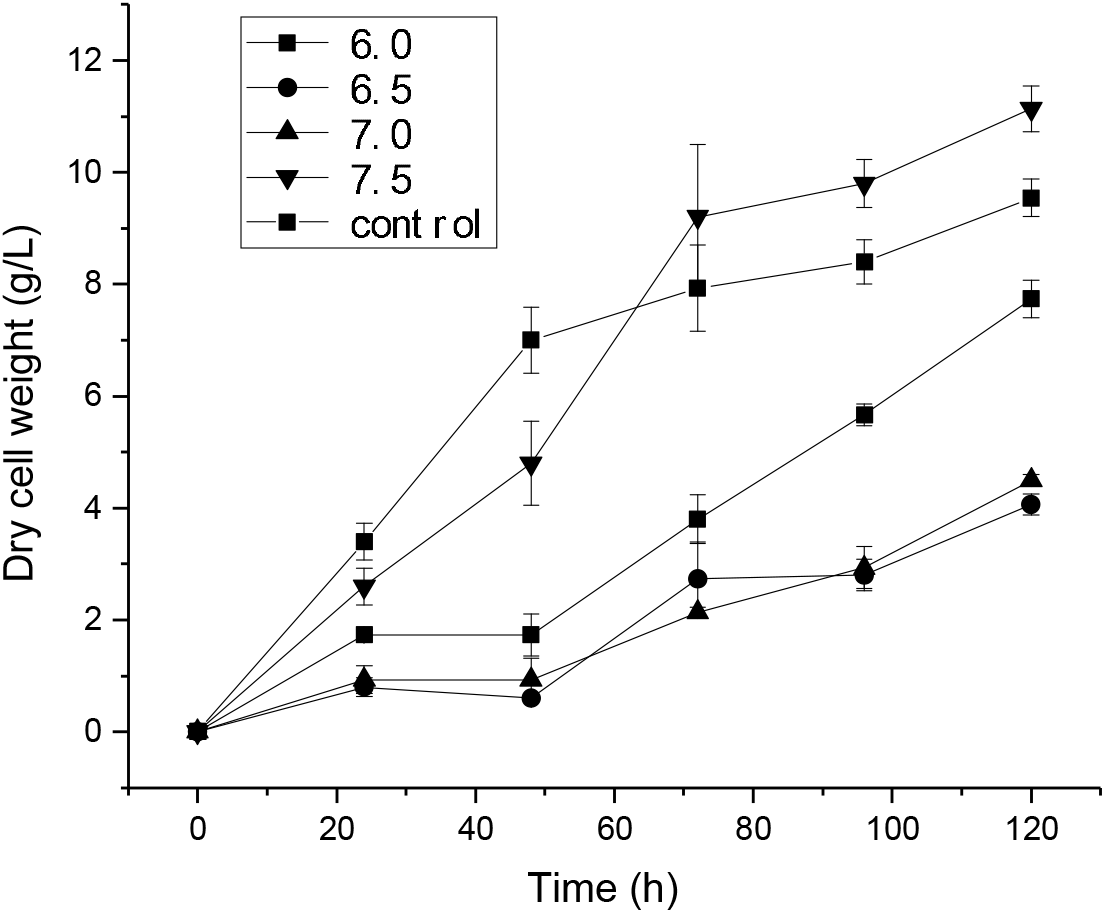
Growth curves of *Y. lipolytica* XP1 in shake flask fermentation with pH 6.0, 6.5, 7.0, and 7.5, the unbuffered media was set as control. All cells were cultivated in CSM-leu media at 30 °C.

Despite the fact that cell growth was rapid under pH 7.5, violacein production did not increase accordingly (Fig. 12). As the media pH was buffered from 6.0 to 7.0, violacein production decreases gradually (Fig. 12), possibly due to the acidophilic nature of the violacein pathway. Most strikingly, CaCO_3_ was found to be a superior supplement to buffer media pH. The carbonate group (CO_3_^2−^) will effectively neutralize the organic acids accumulated during the fermentation. At the end of the fermentation, 70.04 mg/L violacein and 5.28 mg/L deoxyviolacein were produced from the yeast culture supplemented with CaCO_3_, with violacein purity up to 86.92%. This is the highest violacein yield from yeast culture. Due to the inexpensive nature, this result also indicate that CaCO_3_ was the preferred buffer to adjust media pH for yeast fermentation.

**Fig. 12.**
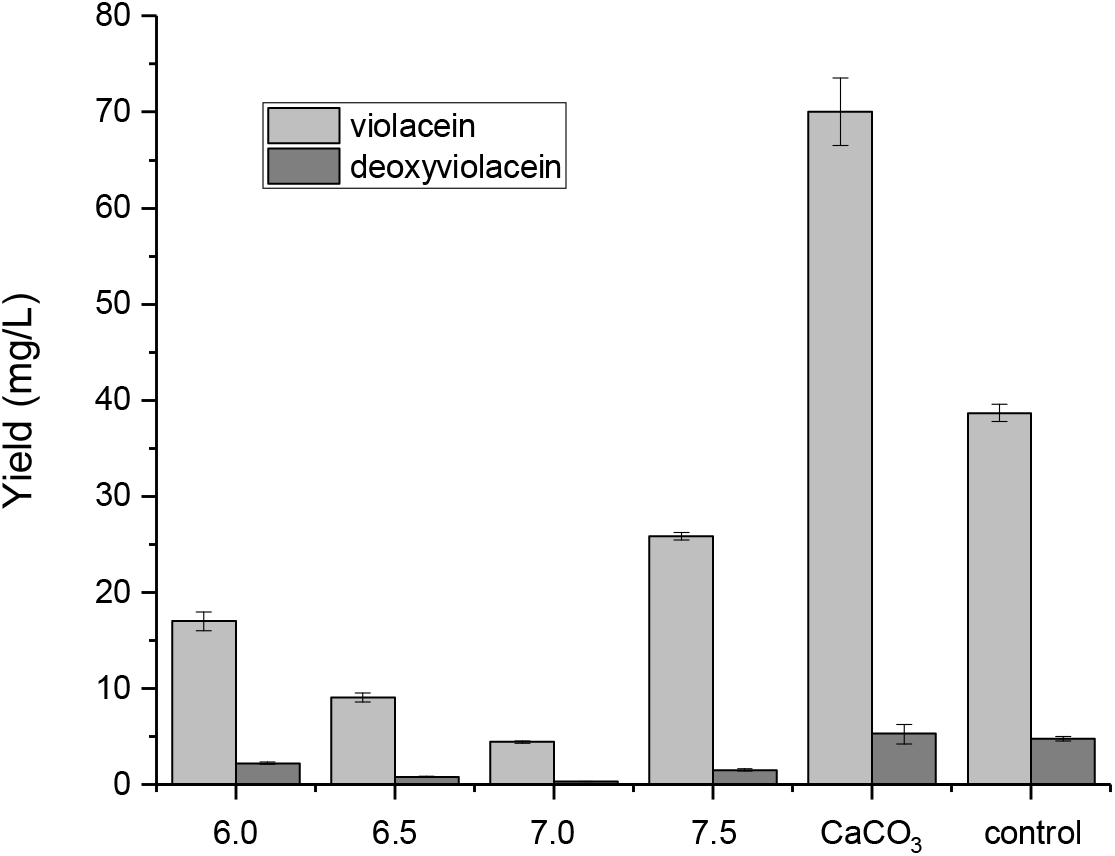
Violacein and deoxyviolacein yield in shake flack fermentation by strain *Y. lipolytica* XP1 with pH 6.0, 6.5, 7.0, 7.5, CaCO_3_ and the control, cultivated in CSM-leu media at 30 °C. Violacein and deoxyviolacein were quantified by HPLC.

## 4 Conclusion

*Y. lipolytica* is considered to be a promising microbial workhorse for production of high value-added products [39, 40]. In this study, we reported the development of quantitative measurement of violacein, based on both HPLC and microplate reader. We demonstrated that both methods are technically equivalent to detect violacein, with the plate reader more suitable for high through-put screening. Due to the rigid cell wall structure, we evaluated a number of extraction procedures to improve the violacein recovery ratio and purity, including the variations of organic solvents, the choice of mechanical shear stress, incubation time and the use of cell wall-degrading enzymes. We found that the highest violacein extraction efficiency could be achieved by adding equal volume of ethyl acetate and grinding with glass beads for 5 minutes. Subsequently, we screened four violacein-producing colonies, optimized the C/N ratio and pH conditions of the yeast fermentation. We found that highest violacein at70.04 mg/L with 86.92% purity could be achieved at a C/N ratio of 60 supplemented with 10 g/L CaCO_3_. This work is the highest violacein production reported in yeast fermentation, and is also the first attempt to optimize the culture condition for violacein production in *Y. lipolytica*. The results reported in this study may be used as a reference for subsequent researches.

## Supporting information

Supplemental data fro Fig. 8

## Acknowledgements

This work was supported by the Cellular & Biochem Engineering Program of the National Science Foundation under grant no.1805139. The authors would also like to acknowledge the Department of Chemical, Biochemical and Environmental Engineering at University of Maryland Baltimore County for funding support. YJT would like to thank the China Scholarship Council for funding support.

## Author contributions

PX conceived the topic. YJT performed analytical assay, extraction and fermentation experiments. YJT and PX wrote the manuscript. YJT was co-advised by JWZ and LZ.

