## Supplemental data fro Fig. 8 for "Engineering oleaginous yeast *Yarrowia lipolytica* for violacein production: extraction, quantitative measurement and culture optimization"

### Supplementary HPLC data for Fig. 8: Violacein production for four biologically-replicated yeast colonies.

| Time(h) | 24 | 48 | 72 | 96 | 120 | 144 | 144 with beads |
|---|---|---|---|---|---|---|---|
| colony 1 | 570.9 | 923.7 | 892.9 | | 892.7 | 951.8 | 987.1 |
| colony 3 | 591.1 | 1003.2 | 997.9 | 1035.8 | 1007.6 | 1069.3 | 1185.4 |
| colony 4 | 605.5 | 899.5 | 830.4 | 801.5 | 879.4 | 949.1 | 1030.2 |
| colony 6 | 550.5 | 976.7 | 962.7 | 930.4 | 950.8 | 1016.5 | 1078.1 |
| | 67.4 | 106.9 | 144.4 | | 273.3 | 213.7 | 536.3 |
| | 48.9 | 120.6 | 174.6 | 271.7 | 351.5 | 301.7 | 762.5 |
| | 49.7 | 105.7 | 125.6 | 195.7 | 208 | 193.8 | 513.1 |
| | 46.3 | 125.2 | 164.7 | 210.2 | 272 | 240.7 | 609.3 |
| | | | | | | | 312.6 |
| | | | | | | | 557.2 |
| | | | | | | | 239 |
| | | | | | | | 384 |

| Time(h) | 24 | 48 | 72 | 96 | 120 | 144 | 144 with beads |
|---|---|---|---|---|---|---|---|
| colony 1 | 6.4279 | 12.4255 | 11.9019 | | 11.8985 | 12.9032 | 13.5033 |
| colony 3 | 6.7713 | 13.777 | 13.6869 | 14.3312 | 13.8518 | 14.9007 | 16.8744 |
| colony 4 | 7.0161 | 12.0141 | 10.8394 | 10.3481 | 11.6724 | 12.8573 | 14.236 |
| colony 6 | 6.0811 | 13.3265 | 13.0885 | 12.5394 | 12.8862 | 14.0031 | 15.0503 |
| | -2.1316 | -1.4601 | -0.8226 | -3.2774 | 1.3687 | 0.3555 | 5.8397 |
| | -2.4461 | -1.2272 | -0.3092 | 1.3415 | 2.6981 | 1.8515 | 9.6851 |
| | -2.4325 | -1.4805 | -1.1422 | 0.0495 | 0.2586 | 0.0172 | 5.4453 |
| | -2.4903 | -1.149 | -0.4775 | 0.296 | 1.3466 | 0.8145 | 7.0807 |
| | | | | | | | 2.0368 |
| | | | | | | | 6.195 |
| | | | | | | | 0.7856 |
| | | | | | | | 3.2506 |

| Time(h) | 24 | 48 | 72 | 96 | 120 | 144 | 144 with beads |
|---|---|---|---|---|---|---|---|
| colony 1 | 4.2963 | 10.9654 | 11.0793 | 14 | 13.2672 | 13.2587 | 21.3798 |
| colony 3 | 4.3252 | 12.5498 | 13.3777 | 15.6727 | 16.5499 | 16.7522 | 32.7545 |
| colony 4 | 4.5836 | 10.5336 | 9.6972 | 10.3976 | 11.931 | 12.8745 | 20.4669 |
| colony 6 | 3.5908 | 12.1775 | 12.611 | 12.8354 | 14.2328 | 14.8176 | 25.3816 |
